# Opportunistic gut sampling indicates differential diet and plastic ingestion risk in Indian dugongs

**DOI:** 10.1101/2021.08.11.455764

**Authors:** Sumit Prajapati, Chinmaya Ghanekar, Sameeha Pathan, Rukmini Shekar, K. Madhu Magesh, Swapnali Gole, Srabani Bose, Sweta Iyer, Anant Pande, Jeyaraj Antony Johnson, Kuppusamy Sivakumar

**Author notes:** **Correspondence** Dr. K. Sivakumar, Scientist-F, Department of Endangered Species Management, Wildlife Institute of India, Chandrabani, Dehra Dun-248001, **Email:**.

## Abstract

Dugongs, exclusively seagrass foragers, are globally threatened marine mammals. Knowledge on their feeding biology has been derived from few direct observations and mostly by analysis of stomach contents. Given limitations in data from Indian populations, dugong strandings serve as an opportunity to understand their dietary composition through gut sampling. In this paper, we utilize the gut contents collected from stranded dugongs to detect differences in the seagrass foraging between two isolated pockets of dugong distribution (Tamil Nadu and Gujarat) and supplement existing knowledge on dugong feeding biology in Indian waters. We extracted, enumerated and identified seagrass species from dugong gut contents. The proportion of seagrass leaf fragments were found higher (>40%) than other fragments in all the gut samples analysed. We recorded two seagrass genera (*Halophila* spp. and *Halodule* spp.) from Gujarat and five seagrass genera (*Halophila* spp., *Halodule* spp., *Cymodocea* spp., *Enhalus* spp., *Syringodium* spp.) from Tamil Nadu dugong individuals. We also obtained anthropogenic debris such as plastic, fishing net and wood fragments from the gut samples. We suggest enhanced monitoring of seagrass habitats and fine spatial scale threat mapping in entire dugong distribution range in India.

---

Dugongs (*Dugong dugon*, Müller, 1776, order Sirenia) are globally threatened marine mammals which primarily forage upon seagrasses (Heinsohn & Birch, 1972; Marsh et al., 2012). Despite a vast global distribution spanning from the east African coast to Australia (Indo-Pacific Ocean region), their populations are declining due to various human-mediated drivers (Marsh et al., 2012). These drivers include mortalities arising from fishnet entanglement, boat strikes, hunting for meat, seagrass habitat loss due to increased sedimentation and pollution (Marsh & Sobztick, 2019), which push the dugong populations towards localised extinctions (Hines et al., 2005; Pusineri et al., 2013; Srinivas et al., 2020).

Dugongs employ two major feeding techniques viz; cropping and excavation of seagrasses, depending on species morphology and substratum (Wirsing, 2007; Rasheed et al., 2016; Marsh et al., 2018). They crop larger and more robust seagrasses species that grow in coarse sediments such as *Enhalus* and *Thalassia* (Erftemeijer et al., 1993; Nakanishi et al., 2008), while smaller seagrasses species growing in fine sediments such as *Halophila, Halodule, Syringodium* and *Cymodocea*, are primarily excavated (Masini et al., 2001; De longh et al., 2007; Sheppard et al., 2007). Dugongs also feed on floating seagrasses (Silas, 1988), algae (Marsh et al., 1982) and invertebrates (active foraging to fulfil nutritional deficiency; Preen, 1995 or passive ingestion while feeding on seagrasses; Preen, 1992). Since availability of seagrasses is a limiting factor, understanding foraging patterns of dugongs is crucial in mapping their distribution. So far, dugong foraging preferences are known through direct observations of feeding or by analysis of stomach contents (Preen, 1992; Andre et al., 2005; De longh et al., 2007; D’souza et al., 2015). Data available from stomach content analysis has provided most detailed information on feeding habits of dugongs (Preen, 1995) and their energy requirements (Andre et al., 2005).

Indian dugong populations are imperilled due to various threats with an estimated population of less than 300 individuals left in the wild (Pandey et al, 2010; Sivakumar & Nair, 2013). Recent efforts at isolated pockets of their distribution along the Indian coastline (Gulf of Kutch in Gujarat; Gulf of Mannar and Palk Bay in Tamil Nadu; and Andaman and Nicobar Islands) have helped generate some crucial ecological data on Indian dugongs including their distribution, important habitats, genetic diversity and connectivity, threats etc. (Sivakumar & Nair, 2013; D’Souza et al., 2013; Rajpurkar et al., 2021; Srinivas et. al, 2020). Limited studies exist on dugong feeding biology from India (D’Souza et al., 2015; Nair & Mohan, 1975) given the difficulty to observe them in the wild. Thus, stranded dugongs provide an opportunity to understand their dietary composition through gut sampling. In this study, we utilize the gut contents opportunistically collected from stranded dugongs to detect differences in the seagrass foraging between study sites to supplement existing knowledge on dugong feeding biology in Indian waters.

In 2018, we sampled gut contents of dead stranded dugongs from the coasts of Tamil Nadu (3 individuals) and Gujarat (2 individuals) (Figure 1). Strandings were informed by the local dugong volunteer network involving fishers and personnel from the Coastal Security Police and State Forest Department (Table 1). Gut samples were collected from foregut of Gujarat dugongs and from both foregut and hindgut from Tamil Nadu dugongs. Samples were preserved in ethanol at −20 [ C until further processing. Dugong carcasses were sexed, measured and necropsies were conducted based on carcass condition (Table 1).

**Figure 1:**
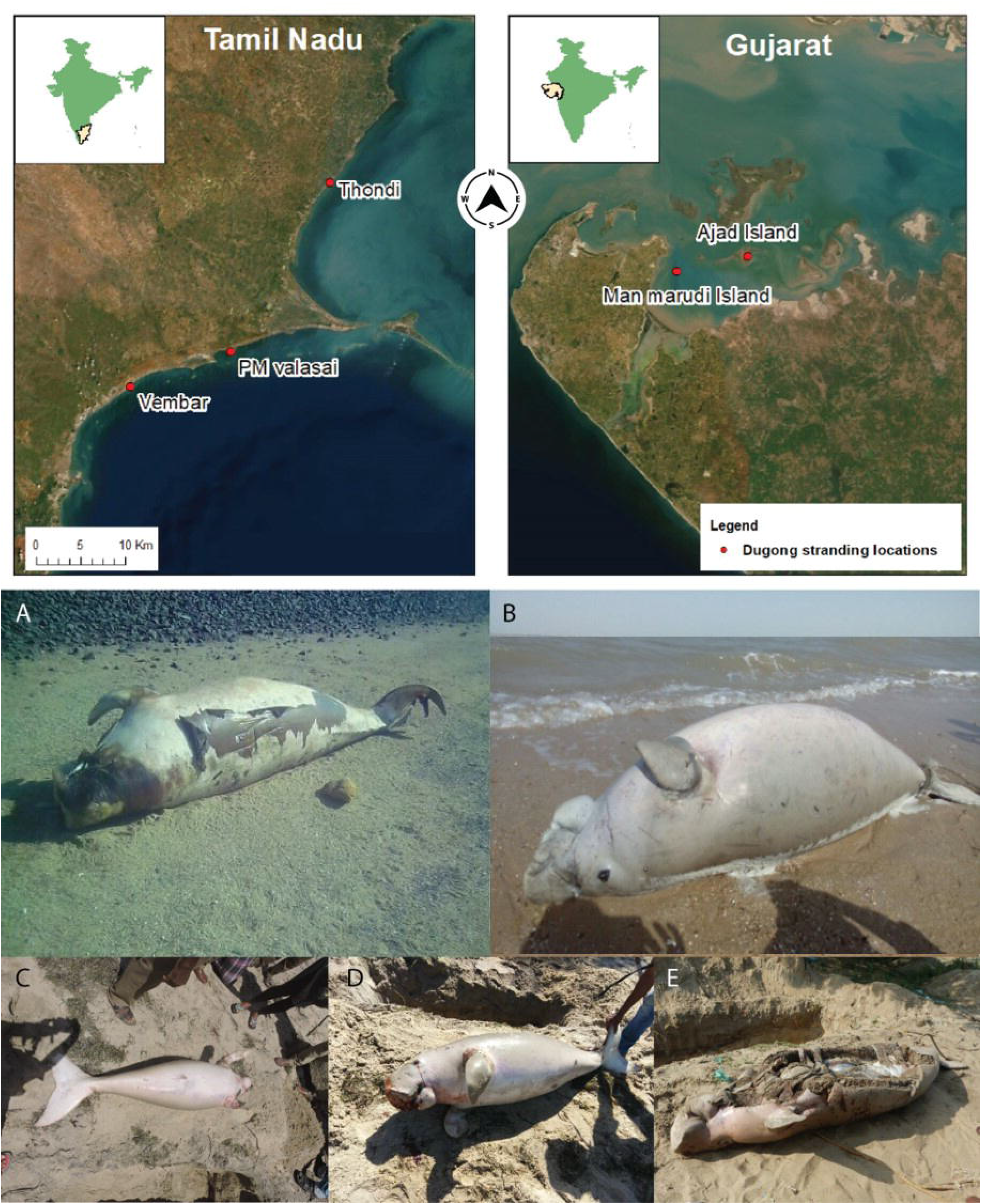
Locations of stranded dugongs in Gulf of Mannar and Palk Bay, Tamil Nadu and Gulf of Kutch, Gujarat. – A) an adult female dugong found in Ajad Island, Gulf of Kutch on 4^th^ February 2018 B) an adult male dugong found in Man Marudi Island, Gulf of Kutch on 20^th^ May 2018 c) juvenile male dugong found in PM Valsai, Palk Bay on 16^th^ June 2018 D) an adult male found in Thondi, Gulf of Mannar on 20^th^ June 2018 E) adult individual found in Vembar, Palk Bay on 7^th^ December 2018

**TABLE 1.** Details on location, carcass condition, cause of mortality and Mean, Percentage contribution of Fragments of Seagrasses and Seagrass genera present in Gut samples of dead stranded dugong from the coast of

| Dugong Individual | Date | Location | Sex | Age class | Carcass condition | Cause of Mortality | Informant group | Body Length (m) | Total gut content weight (g) | Mean ± SD and Percentage proportion of Fragments of Seagrass in /gm samples |  |  | Percentage of Above ground seagrass fragments (leaf and Stem) (%) | Percentage of below ground seagrass fragments (rhizome) (%) | Mean ± SD and Percentage of Fragments (in parenthesis) of Seagrass genera in /gm of Samples |  |  |  |  | Algal Fragments |
| --- | --- | --- | --- | --- | --- | --- | --- | --- | --- | --- | --- | --- | --- | --- | --- | --- | --- | --- | --- | --- |
|  |  |  |  |  |  |  |  |  |  | Leaf | Stem | Rhizome |  |  | *Halophila* spp. | *Halodule* spp. | *Cymodocea* spp. | *Syringodium* spp. | *Enhalus* spp. |  |
| Dugong1 | 04-02-2018 | Ajad Island | Female | Adult | Decomposed | Net Entanglement | Fisherman | 2.6 | 64.77 | 12.13±8.09 (43.29%) | 7.05±3.41 (25.22%) | 8.80±3.80 (31.48%) | 68.51 | 31.48 | 23.8±14.14 (79.20%) | 3.55±4.38 (11.81%) | 0 | 0 | 0 | 2.70±5.18 (8.98%) |
| Dugong2 | 20-05-2018 | Man Marudi Island | Male | Adult | Fresh | Net Entanglement | Fisherman | 2.4 | 159.03 | 14.2±9.15 (40.86%) | 9.30±5.96 (26.76%) | 11.25±6.06 (32.37%) | 67.62 | 32.37 | 4.2±4.81 (6.28%) | 53.65±35.46 (85.36%) | 0 | 0 | 0 | 5.00±5.12 (7.95%) |
| n=2 (Gujarat) | - | - | - | - | - | - | - | - | - | 13.15±8.59 (52.44) | 8.17±5.76 (22.08%) | 10.02±5.15 (25.47%) | 74.52 | 25.47 | 14±14.39 (30.20%) | 25.8±35.67 (61.48%) | 0 | 0 | 0 | 3.85±5.22 (8.30%) |
| Dugong3 | 16-06-2018 | PM Valsai | Male | Juvenile | Fresh | Boat Strike | Fisherman | 1.5 | 76.832 | 7.45±7.43 (41.45%) | 4.60±4.58 (25.59%) | 5.92±3.77 (32.94%) | 67.04 | 32.94 | 8.82±9.21 (44.08%) | 5.72±6.85 (28.59%) | 1.02±7.56 (5.10%) | 4.35±6.25 (21.74%) | 0.1±0.30 (0.50) | 0 |
| Dugong4 | 20-06-2018 | Thondi | Male | Adult | Fresh | Poached | Forest Department and Marine Police | 2.4 | 25.94 | 16.55±1.0 (55.29%) | 8.05±6.46 (26.89%) | 5.30±2.55 (17.80%) | 82.19 | 17.80 | 16.45±6.87 (49.76%) | 1.86±1.46 (5.63%) | 2.50±1.57 (7.56%) | 12.08±8.51 (36.45%) | 01.±0.30 (0.30) | 0.1±0.30 (0.30%) |
| Dugone5 | 07-12-2018 | Vembar | Not identified | Adult | Decomposed | Na | Fisherman | 2.9 | 30.73 | 21.65±7.31 (60.81%) | 5.25±4.38 (14.74%) | 8.70±2.61 (24.43%) | 75.55 | 24.43 | 5.50±2.96 (7.68%) | 4.15±2.47 (5.80%) | 61.8±14.14 (86.31%) | 0 | 0 | 0 |
| n=3 (Tamil Nadu) | - | - | - | - | - | - | - | - | - | 13.30±10.27 (41.95%) | 5.60±5.20 (26.06%) | 6.46±3.46 (31.97%) | 68.01 | 31.97 | 9.50±4.89 (26.49%) | 4.36±5.26 (12.16%) | 16.58±27.31 (46.24%) | 5.32±7.45 (14.83%) | 0.07±0.26 (0.19) | 0.025±0.15 (0.069%) |

We categorized seagrass fragments from the gut samples as leaf, stem and rhizome based on their morphological features, to understand the proportion of above ground (leaf, stem) and below ground (rhizome) plant material (Figure 2) (Adulyanukosol & Poovachiranon, 2003). We used point-intercept method (Schuette et al., 1988) to calculate the abundance of fragments of leaves, rhizomes and stems. Samples were further divided into 10 subsamples of 1 gram each and later homogeneously spread on petri plate of size 100 × 15 mm. The petri plate was divided into 24 quadrates of 1cm^2^ each and the gut material was observed under stereo-microscope (10x magnification). We identified seagrass leaf fragments based on apex structure, visible venation, stem fragments from its fibroid structure and rhizomes were identified by their hard structure with nodes and presence of leaf scars (Adulyanukosol & Poovachiranon, 2003).

**Figure 2:**
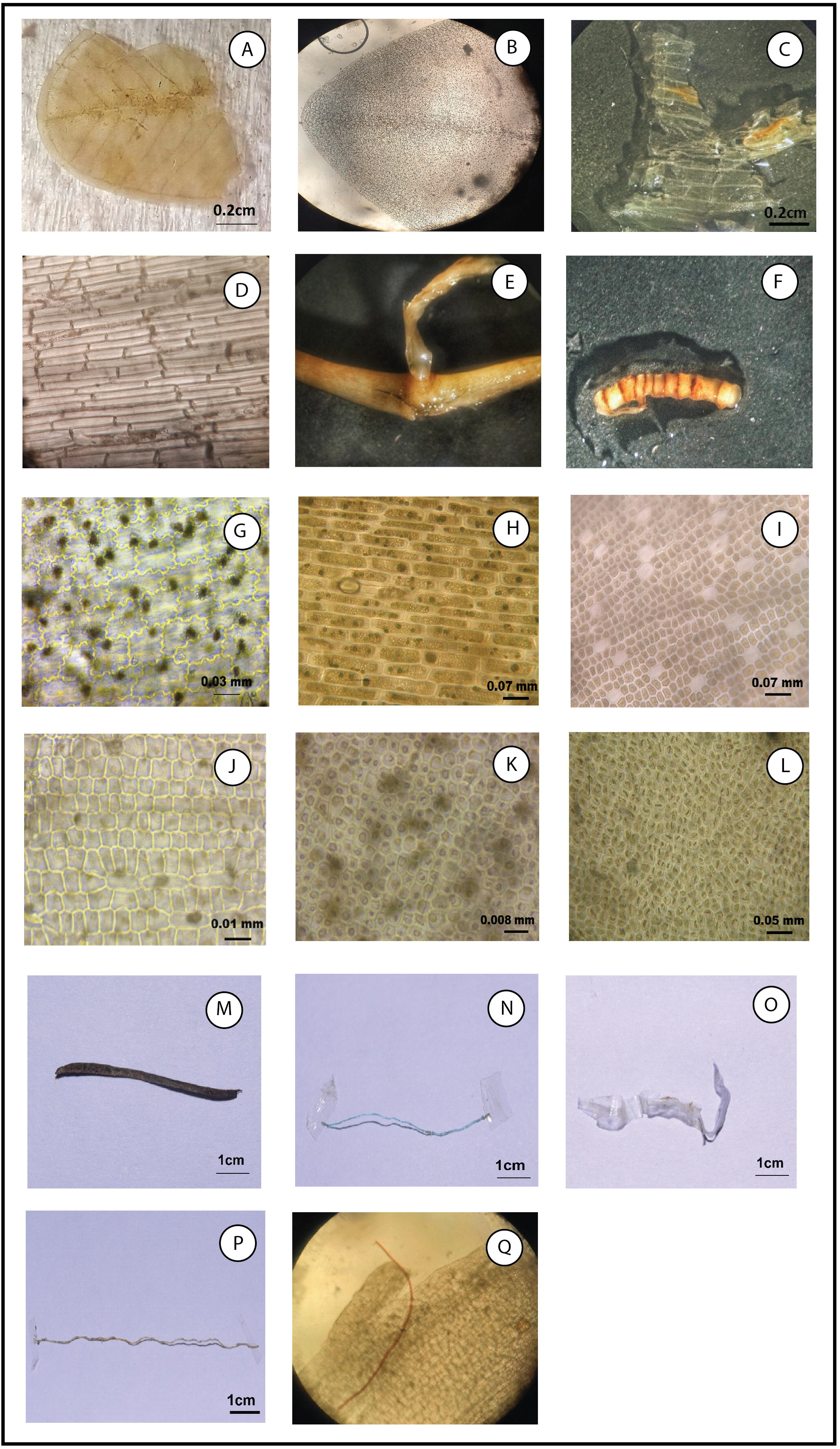
Fragments and epidermal cell structures of seagrasses and non-biological materials from dugong gut samples – A) & B) Leaf fragment of *Halophila* spp. with venation under stereo-microscope and compound microscope (4x); C) fibroid structure of vertical stem under stereo-microscope; D) epidermal cell structure of stem under compound microscope (20x); E) rhizome fragment; F) Presence of leaf scars in rhizome fragment under stereo-microscope; G) epidermal cell structure of *Halophila* spp. (40X); H) epidermal cell structure of *Halodule* spp. (40x); I) epidermal cell structure of *Cymodocea* spp. (40x); J) epidermal cell structure of *Enhalus* spp. (40X); K) epidermal cell structure of *Syringodium* spp. (40X); L) epidermal cell structure of an algal fragment (40X); M) wooden fragment (4cm); N) fishing net filament (4cm); O) polythene fragment (4cm); P) fishing net fragment (9cm); Q) red-coloured microfilament (20X)

We counted leaf fragments for seagrass genera using the quadrat method as stem and rhizome are conserved in appearance through most genera (Channells & Morrissey, 1981). One-quarter of the petri plate (25% grids) was chosen randomly to count the leaf fragments. Epidermal cell characteristics and tannin cell arrangements were used to identify the seagrass up to genera level for macerated samples (Channells & Morrissey, 1981). Gross morphological features (venation, size, apex structure and shape of leaves) were taken into consideration for intact leaves (Adulyanukosol & Poovachiranon, 2003). Fragments were identified at different magnifications (4x, 10x and 40x) using seagrasses identification keys (Lanyon, 1986; El Shaffai, 2016; Pande et al., 2021). Species level identification was not possible as epidermal cell structures are similar for two species of the same seagrass genus (Adulyanukosol & Poovachiranon, 2003). Mann-Whitney *U* test was performed for *Halophila* spp. and *Halodule* spp. to check differences in their occurrence between sites i.e., Tamil Nadu and Gujarat as well as between fresh and decomposed carcass. This test was possible only for these two genera, as these occurred in all the gut samples.

We also collected eight species of seagrasses from Tamil Nadu, Gujarat and Andaman & Nicobar Islands. Morphology of leaf and epidermal cells’ features were studied and photographed under stereo (10x) and inverted microscopes in various magnifications (4x, 10x, 20x and 40x). These slides were used as reference for seagrass identification, up to genera level. Identification of seagrass genera found in gut samples were reconfirmed with epidermal cells of reference seagrass samples collected from a fore mentioned sites field sites see (Supplementary Figure 1).

The proportion of seagrass leaf fragments were found higher (>40%) than other fragments in all the gut samples analyzed (Table 1). In the gut content of Gujarat individuals, the percentage of above ground fragments (leaf, stem) was double the underground plant fragments, whereas one sample individual from Tamil Nadu exhibited similar proportions (Table 1). Two of three samples from Tamil Nadu showed higher proportions of above ground fragments than the below ground (Table 1). We recorded two seagrass genera from Gujarat dugong individuals, *Halophila* spp. and *Halodule* spp. (Figure 2). *Halodule* spp. (61.48 %) fragments were more abundant than *Halophila* spp. (30.20 % with little algal fragments (8.30 %) (Table 1). Five seagrass genera namely *Halophila* spp., *Halodule* spp., *Cymodocea* spp., *Enhalus* spp., *Syringodium* spp. and alga were recorded from Tamil Nadu dugong individuals (Figure 2). Leaf fragments of *Cymodocea* spp. (46.24%) were dominant followed by *Halophila* spp. (26.49 %), *Syringodium* spp. (14.83 %), and *Halodule* spp. (12.16%). Low occurrence of *Enhalus* spp. (0.19 %) and algal fragments (0.069 %) were found in the samples (Table 1). We found *Halodule* spp. occurrence to differ between Tamil Nadu and Gujarat samples (U=987.5, *p* < 0.001) but not for *Halophila* spp. (U=1411, *p* > 0.05; Figure 3). Occurrence of *Halophila* spp. was observed to differ according to carcass condition (U=1214, p<0.05) but not *Halodule* spp. (U=1840, p=>0.05; Figure 3).

**Figure 3:**
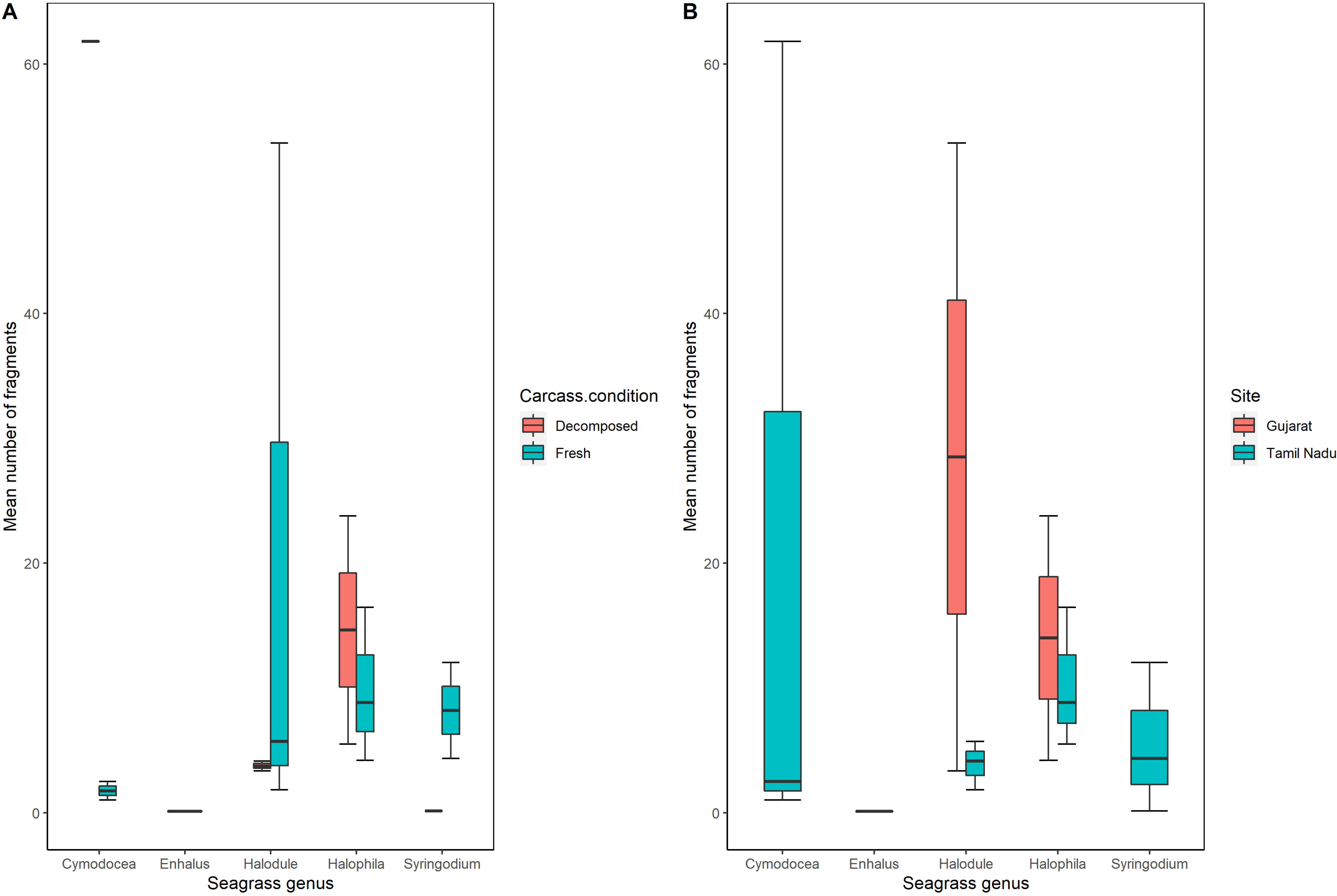
Mean number of seagrass fragments obtained from the gut of stranded dugongs sorted according to A) carcass condition and B) sampling site.

In addition to seagrasses, non-plant materials i.e., plastic and wooden fragments (Figure 2) were found in the gut content of two individuals (one each from Gujarat and Tamil Nadu). Two fishing net filaments (~9.4 cm and ~ 4 cm in length), one polythene fragment (~4cm length) and one wooden fragment (~4cm in length) were obtained from Tamil Nadu dugong, while one plastic microfilament was retrieved from Gujarat dugong (Figure 2).

Globally, studies on dugong gut content have highlighted selective consumption of seagrass species like *Halophila ovalis, Halodule uninervis* (Heinsohn & Birch, 1972; Johnstone & Hudson, 1981; Aragones 1994, Adulyanukosol et al., 2010), *Enhalus* spp. (Erftemeijer et al., 1993), *Thalassia hemprichii, Syringodium isoetifolium* (Andre et al., 2005) and *Cymodocea serrulata* (Andre et al., 2005; Adulyanukosol et al., 2010). Percentage contribution of above and below ground plant material is proportionate in the gut samples of Gujarat individuals (Table 1) which suggests towards dugongs excavating the whole seagrass plant. In Tamil Nadu, two of the three samples exhibited higher preference of cropping mechanism over excavation (% of above ground seagrass fragments > % of below ground seagrass fragments) (Table 1), which may result from higher disturbance rate from the concerned study area. We need more ecological observations and larger sample size, supported with threat analysis to validate from our study sites. Limited percentage of algal material (3.58%) could be due to incidental ingestion while feeding on seagrasses. In the present study, more generic diversity of seagrasses in gut content of individuals from Tamil Nadu (n=5) than Gujarat (n=2), could be attributed to high regional generic diversity of seagrasses in Tamil Nadu (Thangaradjou & Bhatt, 2018) than that of Gujarat (Thangaradjou & Bhatt, 2018).

Dominance of *Halophila* spp. in dugong gut content samples from Ajad Island, Gujarat is in line with field observations of seagrass habitats in the region which is dominated by meadows of *H. ovalis* and *H. beccarii* (Sivakumar et al., 2020), strongly suggesting this region to be potential dugong foraging habitat. *Halodule* spp. was found to be dominant in the samples of Man Marudi Island (close to Beyt Dwarka), Gujarat where *Halophila ovalis* and *Halophila decipiens* meadows are more common in occurrence (Sivakumar et al., 2020). Considering a low population size of dugongs in Gulf of Kutch (Pandey et al, 2010), locating *Halodule* spp. meadows would help in identifying critical foraging grounds in the area.

Differential digestion rates of seagrass species is known to change their relative abundance in the digestive tract of dugongs (Thayer et al., 1984; Preen, 1995); which could further be affected by carcass’ condition and decomposition rate of individual species. Additionally, role of plant morphology and composition are also crucial factors affecting time required for digestion and in turn occurrence in the gut (Heinsohn & Birch., 1972). More fibrous species like *Cymodocea* spp. take longer time for decomposition (Marsh et al., 2018) and digestion as compared to smaller leaved, less fibroid plants like *Halophila* spp. Difference in *Halophila* spp. fragments in both fresh and decomposed carcasses can be attributed to its availability and differential digestion rate (Lanyon & Sanson, 2006).

Stranded dugongs in Tamil Nadu shows a differential rate of decomposition of seagrass diet from fresh carcasses in comparison to highly decomposed state (Table 1). Fresh carcasses were recorded with higher mean occurrence of *Halophila* spp., while decomposed carcasses were diagnosed with *Cymodocea* spp. in dominance. This could be attributed to carcass condition herein all seagrass genera (low and high fibre) were present carcasses while only high fibroid genus like *Cymodocea* spp. were retrieved from the decomposed one. Less proportion of *Enhalus* spp. in two carcasses and total absence from the third one (Table 1), which suggests incidental feeding on the species.

Overall, this study generates key bassline data on dugongs from two of its important distribution regions in the Indian sub-continent. Occurrence of plastic, fishing net fragment and wood debris indicates towards a potential risk to dugongs at their forage grounds. Thus, we recommend ramping up monitoring of seagrass habitats and fine spatial scale threat mapping in entire dugong distribution range in India.

## ACKNOWLEDGEMENTS

This study was sponsored by National CAMPA Advisory Council (NCAC), Ministry of Environment, Forest and Climate Change, Government of India (Grant/Award Number: 13-28(01)/2015-CAMPA). We acknowledge the state forest departments of Tamil Nadu and Gujarat for providing logistics support at field sites. We thank the director, dean, research coordinator, and nodal officer (external projects) of the Wildlife Institute of India for their constant support. We also thank Vabesh Tripura, Sohom Seal and Ankita Anand for help in preparation of figures. We are grateful for the generous field support of the fisherfolk and volunteers at field sites.

---

**Supplementary Figure 1:** Enlarged images (40X) of epidermal cells of seagrass species collected as reference material. A) *Halophil*a spp. B) *Halodule* spp. C) *Cymodocea* spp. D) *Enhalus* spp.

