## Supplementary figures and images for "Opportunistic gut sampling indicates differential diet and plastic ingestion risk in Indian dugongs"

### Supplementary Figure 1

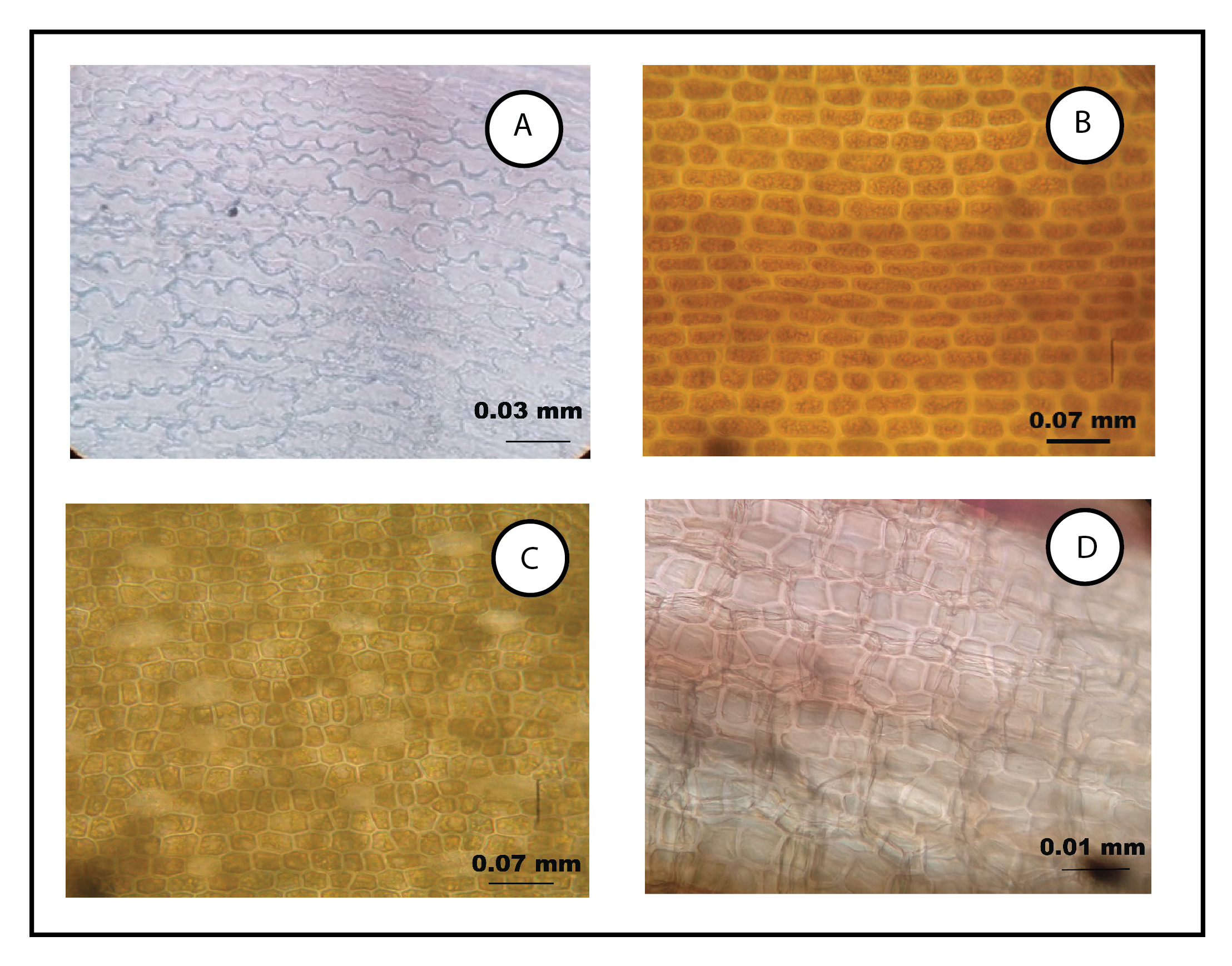
